# Low-level carbon monoxide exposure affects BOLD FMRI

**DOI:** 10.1101/141093

**Authors:** Caroline Bendell, Shakeeb H. Moosavi, Mari Herigstad

## Abstract

Blood Oxygen Level Dependent (BOLD) FMRI is a common technique for measuring brain activation that could be affected by low-level carbon monoxide (CO) exposure from e.g. smoking. This study aimed to probe the vulnerability of BOLD FMRI to CO and determine whether it constitutes a significant confound in neuroimaging and clinical trials. Low-level (6ppm exhaled) CO effects on BOLD signal were assessed in 12 healthy never-smokers on two separate experimental days (CO and air control). FMRI tasks were breath-holds (hypercapnia), visual stimulation and fingertapping. CO significantly dampened global BOLD FMRI signal during hypercapnia and visual cortex activation during visual stimulation. During fingertapping, CO reduced visual cortex activation but increased premotor cortex activation. Behavioural and physiological measures remained unchanged. We conclude that BOLD FMRI is vulnerable to CO, possibly through baseline increases in CBF, and suggest exercising caution when imaging populations exposed to elevated CO levels, e.g. with high smoking prevalence.

## INTRODUCTION

One of the most common methods used to measure brain function in humans is FMRI, of which Blood Oxygen Level Dependent (BOLD) FMRI is arguably the most mainstream technique. BOLD FMRI is an indirect measure of brain activation, based on changes in the ratio of oxygenated to deoxygenated blood in the brain, which depends on cerebral metabolic rate (CMRO_2_), cerebral blood volume (CBV) and cerebral blood flow (CBF). These factors may be altered as part of the experimental design or as unintended confounds, potentially affecting BOLD signal.

Carbon monoxide (CO) is a cerebral vasodilator (1, 2). Increases in CBF with elevations in CO has been shown in animal models (2-6) as well as in humans (7, 8). Low-level CO exposure is common, in particular through inhalation of cigarette smoke. Smokers typically have elevated carboxyhaemoglobin (COHb) levels (6ppm in exhaled air or above (9)) compared to nonsmokers (1-5ppm). As smoking behaviour is associated with e.g. socioeconomic status and disease status, elevated COHb may significantly influence neuroimaging results on the group level in such demographics. If CO causes a baseline increase in CBF, the capacity for further local vasodilation and thus detectable BOLD signal could be reduced. As the BOLD signal for any given task is assessed by comparing task-related signal to baseline signal, this altered baseline could artificially dampen the observed task-specific BOLD response.

One method to test cerebral vasodilation is carbon dioxide (CO_2_) exposure. Hypercapnia induces a strong CBF increase but no change in oxygen metabolism (10). It has been used as a cerebrovascular challenge in FMRI studies (11-13) due to its global and reproducible effect on BOLD signal (12). Increased baseline CBF can reduce the vascular responsiveness to hypercapnia, and as the BOLD response to CO_2_ is strong, even a small increase in baseline CBF might have a discernible effect on BOLD signal. Moreover, hypercapnia can be readily introduced through breath holds (accumulating CO_2_ in the blood stream). In this study, we therefore used the hypercapnia derived through breath holds as a tool to investigate the effect of CO on BOLD signal.

We hypothesised that low-level CO inhalation would significantly reduce global BOLD signal during hypercapnia. To determine whether the effect of CO is robust enough to extend to common FMRI paradigms, we also included a simple visual stimulation and motor task, hypothesising that CO would dampen BOLD signal in brain regions associated with these tasks.

## METHODS

### Participants

We recruited 12 (8F, age 25.3 years +/−4.3) healthy never-smokers to the study. Exclusion criteria were MRI contraindications, smoking history, history of cardiorespiratory or neurological disease, and pregnancy (women only). All participants gave written, informed consent. The study was approved by Oxford Brookes University Research Ethics Committee.

### Protocol

Participants were asked to attend a preliminary laboratory visit. During this visit, medical history and state and trait anxiety inventory (STAI) questionnaires were completed (14). A breathing test was conducted to let the participant familiarise themselves with the breathing system and the CO exposure. Participants were asked to breathe on a custom-made breathing system through a mouthpiece with their nose occluded, and were given time for their breathing to stabilise before commencing the experiment. After stable breathing had been recorded for five minutes, CO was added to the inspired air over five minutes, out of sight of the participant. Following CO administration, five more minutes of stable breathing was recorded. During the experiment, ECG, pulse pressure and saturation was continuously measured. Expired CO measurements were made before, immediately after, and 10 minutes after the breathing test (Micro+ Smokelyzer, Intermedical Ltd., Kent).

MRI scans were conducted on two separate days (Figure 1). Participants were asked to complete a state anxiety questionnaire on arrival and after the end of the experiment on each day. Whilst in the scanner, participants were asked to undertake the following tasks: breath holds, a visual stimulation task, a motor task and a simple reaction time task. We chose to investigate a set of commonly used tasks of short duration to investigate whether quantifiable differences could be observed in BOLD response due to CO even when tasks are underpowered. Breath holds were conducted end-expiration, signalled by visual cues (‘Get ready’ two seconds before breath hold, ‘Hold your breath’ during breath hold) and lasted 15 seconds, repeated 4 times with 15 second breaks in between. The visual stimulation was a flashing checkerboard (8Hz, lasting 10 seconds, separated by 10-second breaks, repeated thrice). The motor task was tapping of the right index finger, signalled by visual cues (‘Tap your right index finger’) and lasted 15 seconds, repeated 4 times with 15-second breaks in between. The reaction time task required participants to immediately press a button upon the appearance of a red dot on the screen (24 appearances, random intervals). These tasks were conducted twice, once before the breathing intervention (baseline) and once after (postintervention). On one day, the intervention was air, and on the other day, the intervention was CO. Participants were not aware of which intervention would be given on any of the days and the order of the interventions was randomised and balanced. Training in all FMRI tasks were given by an experimenter prior to the first scan on each day, to ensure that the participant could reliably complete these on their own in the scanner. Expired CO measurements were made before the first scan, immediately after the second scan (~20 minutes after the breathing intervention) and 10 minutes after the second scan (~30 minutes after the breathing intervention).

**Figure 1.**
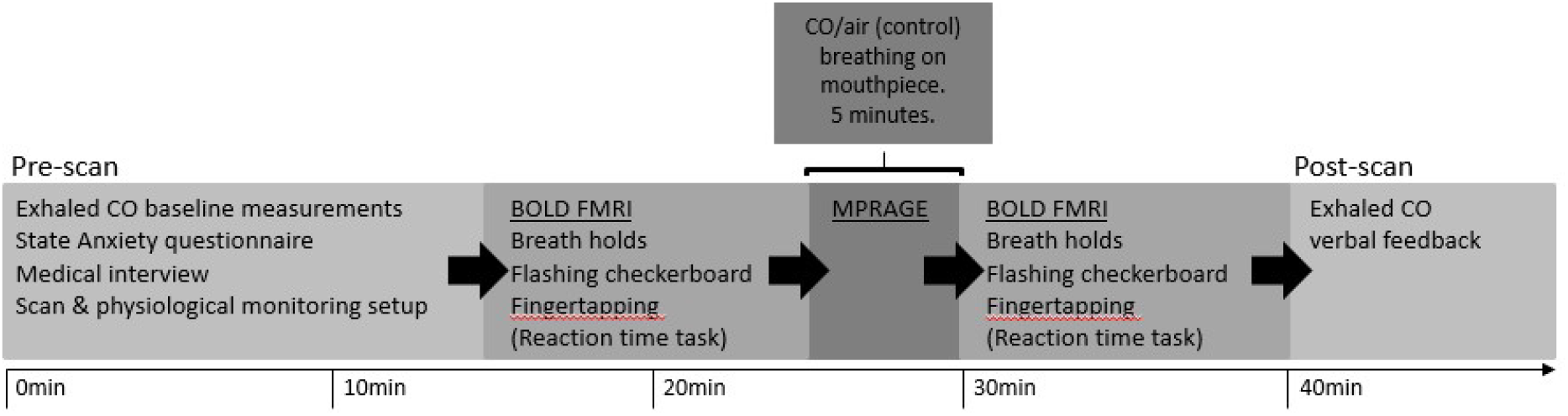
Schematic of protocol. FMRI tasks included breath-holds (end-exhalation), visual stimulation (8Hz flashing checkerboard) and a (right hand) finger-tapping task. Two sets of BOLD scans were obtained on each experimental day, separated by a 5 min breathing intervention (air or CO, order randomized and counterbalanced) during which a structural scan was acquired.

### MRI data acquisition

Imaging was performed at the University of Oxford Centre for Clinical Magnetic Resonance Research with a Siemens 3Tesla TIM-Trio scanner, using a 12-channel head coil. Participants were given two FMRI scans (BOLD echo-planar image acquisition, time repetition (TR) = 3000ms, time echo (TE) = 30ms, field-of-view = 192x192mm, voxel-size = 3x3x3mm, 45 slices) on each day, separated by the intervention period (air or CO). A structural T1-weighted, whole-brain scan (MPRAGE, TR = 2040ms, TE = 4.7ms, flip angle = 8°, voxel-size = 1x1x1mm) was used for image registration.

Heart rate (HR) and pulse oximetry (SaO_2_, multigas monitor, 9500, MR Equipment), ECG, respiration (respiratory bellows around the chest) and end-tidal partial pressures of oxygen (P_ET_O_2_) and CO_2_ (P_ET_CO_2_; Datex, Normocap) were continuously measured throughout the scans. ECG data was observed throughout. All other physiological data were sampled at 50Hz and recorded along with scan volume triggers via PowerLab 16/35 using LabChart (ADinstruments).

### Data Analysis

FMRI data processing was carried out within FSL (Oxford Centre for Functional Magnetic Resonance Imaging of the Brain (FMRIB) Software Library, using FEAT (FMRI Expert Analysis Tool) Version 5.98. The cluster Z threshold was set to 2.3 and a corrected cluster significance threshold to p=0.05.

Prestatistic processing of the data included MCFLIRT motion correction (15), spatial smoothing with a full-width-half-maximum Gaussian kernel of 5mm and high-pass temporal filtering (Gaussian-weighted least-squares straight line fitting, high-pass filter cut-off of 60s). FSL motion outliers was used to detect and regress out large motion artifacts. Data was modelled using FMRIB’s Improved Linear Model (FILM) with local autocorrelation correction (16). Images were registered to the MNI152 standard space using an affine registration between the EPI and T1-weighted scan and a nonlinear registration between the T1-weighted scan and the MNI standard brain.

General Linear Models (GLMs) with multiple explanatory variables (EVs) incorporating timing values for the different events were designed to describe the data. A physiological noise modelling tool was used to regress out effects of physiological noise (17), and CO_2_ measurements added as a separate regressor in the GLM for all tests. A 6-second haemodynamic delay was assumed and contrast images were used for higher-level analyses as appropriate.

A fixed-effects model was used to generate contrast of parameter estimate (COPE) images of the difference between the baseline and post-intervention scans for each participant on each experimental day. This was done by forcing random effects variance to zero in FLAME (FMRIB’s Local Analysis of Mixed Effects) (18, 19). Means of all COPE images were calculated for both protocols. Group analyses compared COPE images between protocols for each task using a whole-brain approach. In the group analyses, contrasts of interest were CO_2_ fluctuations with breath holds, presentation of flashing checkerboards (visual task) and finger-tapping.

STAI questionnaires were scored according to their respective manuals and compared between sessions using nonparametric t-tests (Mann-Whitney U test). Physiological data was analysed using custom-written MATLAB scripts and compared using Student’s t-tests. Data obtained during the motor task was used for comparison of end-tidal gases between protocols.

## RESULTS

Participant details and behavioural data are presented in Table 1, and physiological data is presented in Table 2.

**Table 1.** Participant details and behavioural data

| **Table 1. Participant details and behavioural data** |  |  |  |
| --- | --- | --- | --- |
|  | preliminary visit | MRI visit (CO) | MRI visit (Air) |
| Sex (F/M) | 8/4 | 8/4 | 8/4 |
| Age (years) | 25.3 (4.3) | 25.3 (4.3) | 25.3 (4.3) |
| BMI (kg/m<sup>2</sup>) | 23.6 (3.0) | 23.6 (3.0) | 23.6 (3.0) |
| Trait anxiety score | 35.4 (7.2) [23-47] | N/A | N/A |
| State anxiety score | 31.1 (8.6) [21-55] | 27.0 (4.3) [21-35] | 28.2 (4.4) [23-37] |
| Change in average reaction time (post > pre) | N/A | 15.2 (24.3) | 10.0 (21.8) |

Participant details (n=12), averages +/- standard deviation. Range is provided for anxiety scores. BMI=body mass index

**Table 2.**
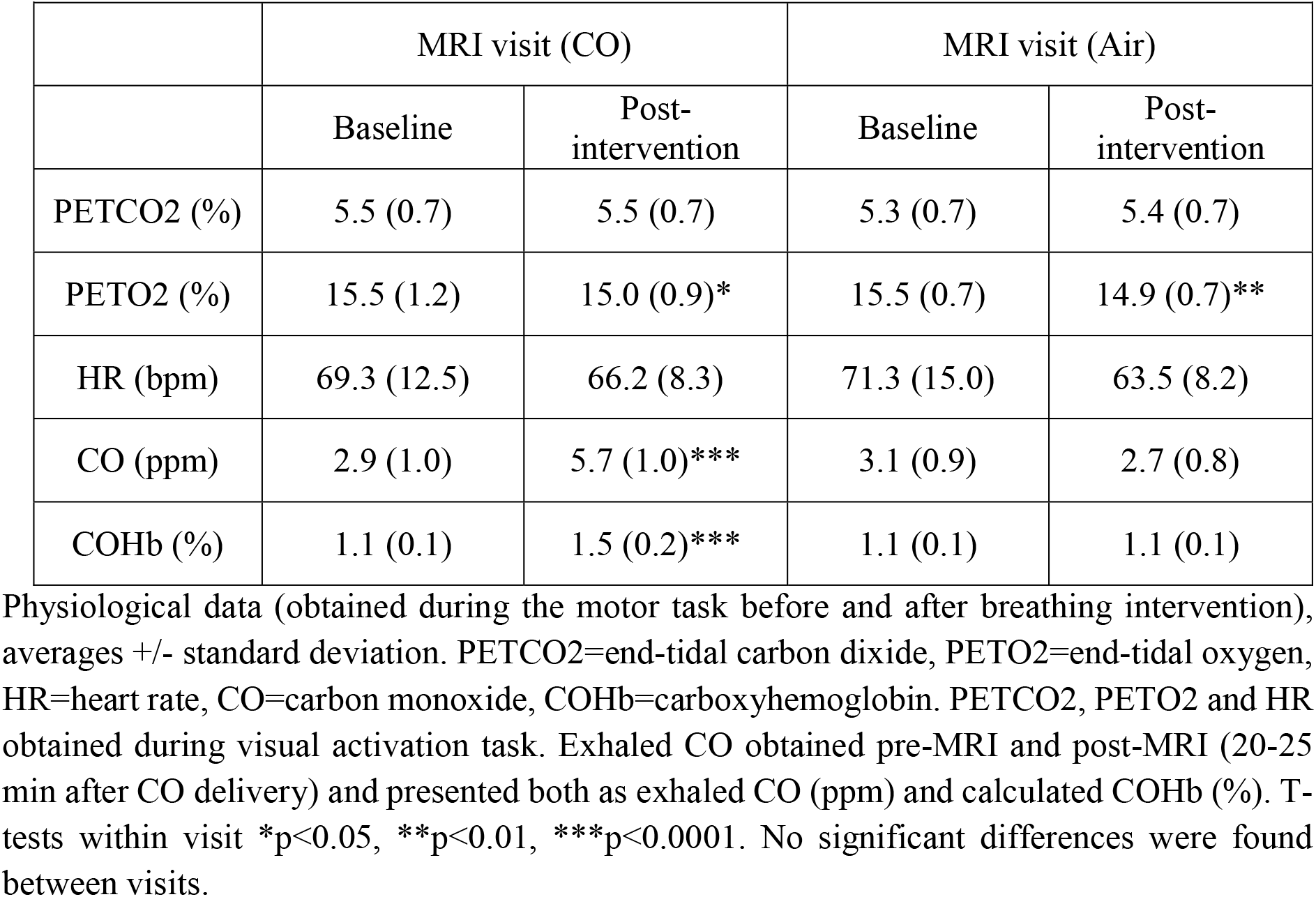
Physiological data

### Psychological and physiological data

There were no significant differences between protocols in anxiety scores (p=0.55) or reaction times (p=0.29, adjusted for baseline scores). None of the participants was able to discern between CO and air inhalations when prompted. P_ET_O_2_ was significantly reduced between baseline and post-intervention scans in both protocols, but no significant difference was found between protocols (p=0.57). There was no change in P_ET_CO_2_ or HR between scans or protocols. CO values increased significantly in the CO protocol (p<0.0001, Figure 2), but not air (p=0.29). CO exposure increased average exhaled CO from 2.9+/−1.0 ppm to 5.7+/−1.0 ppm.

**Figure 2.**
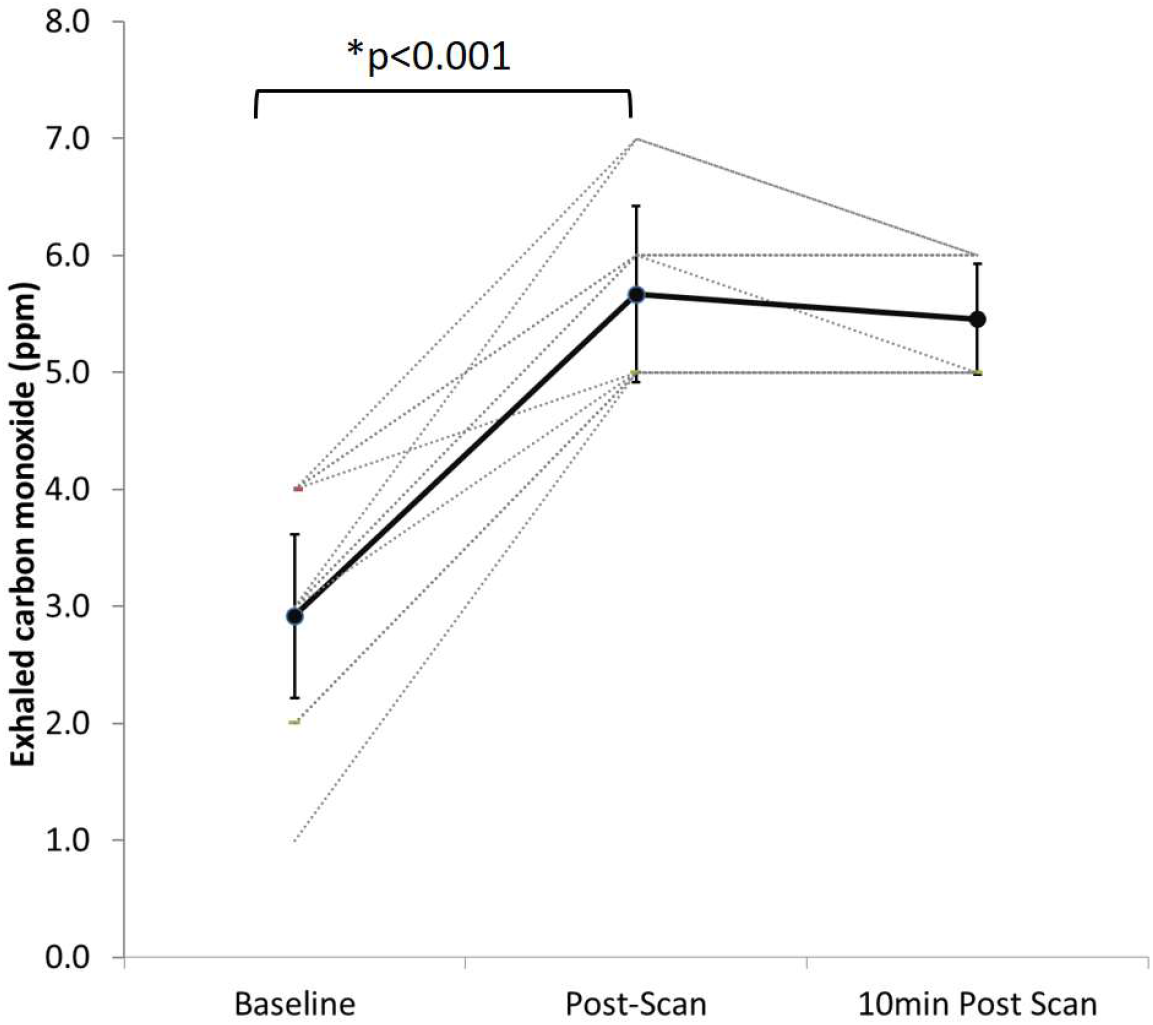
Exhaled CO (ppm). Baseline, post-scan (~20 min after end-inhalation) and 10 min post-scan (~30 min after end-intervention). Individual values plus average and standard deviation (bold line).

### FMRI data

For reported FMRI results, significance denotes thresholded, cluster corrected, activation (p<0.05).

*Breath hold task (Figure 3).* The rise in CO_2_ with breath holds caused signal increase throughout the grey matter during all scans. Mean global signal was markedly lower following the CO intervention compared to baseline scans and after the control (air) intervention. Group comparisons showed reduction in activation in the CO protocol compared to air in the left operculum and insula, left middle and inferior frontal gyri, left dorsolateral prefrontal cortex, left lateral occipital cortex, left supramarginal and angular gyri and the brain stem.

**Figure 3.**
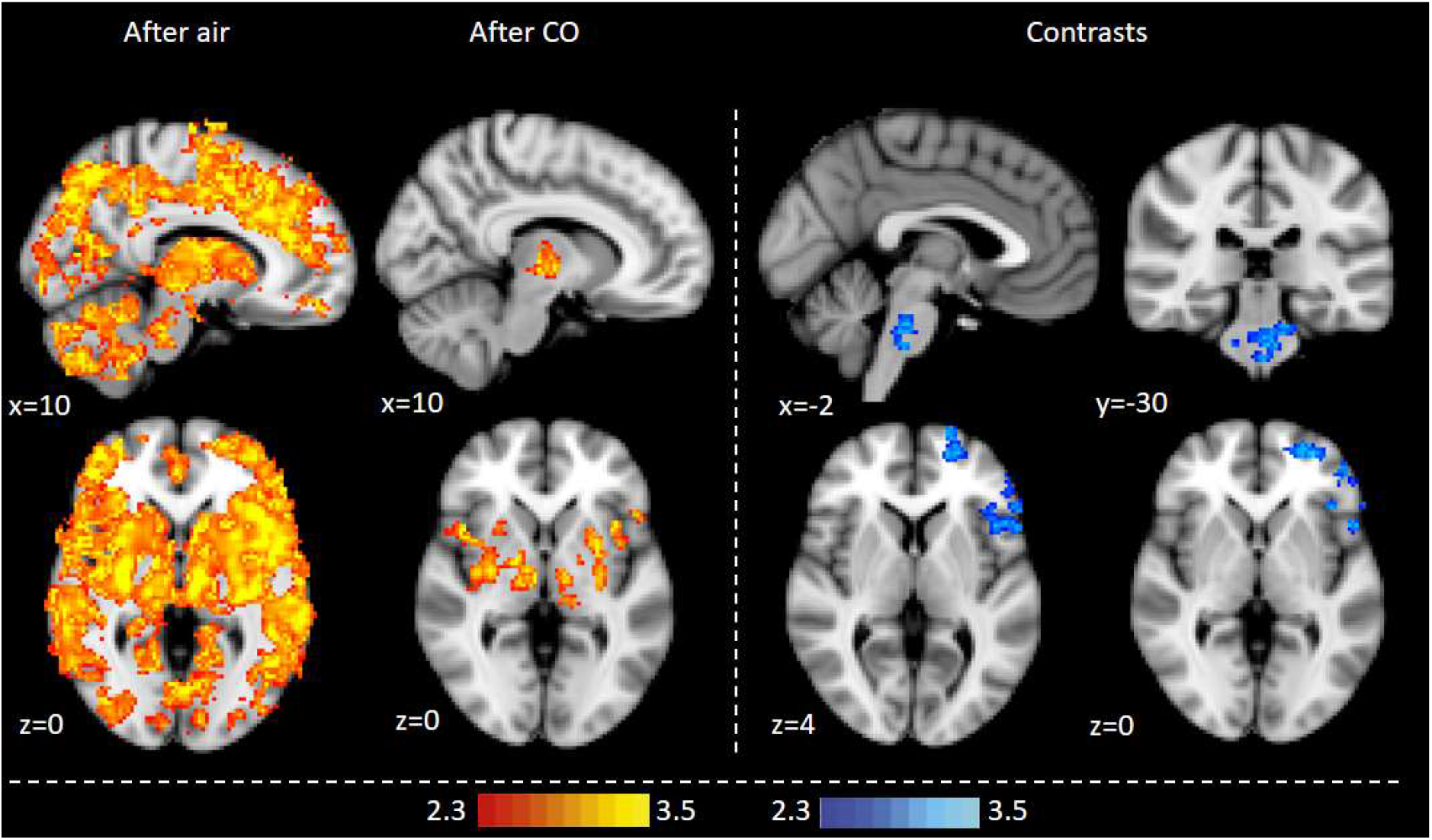
BOLD FMRI activation with CO_2_ variation (breath holds). Whole-brain analysis, cluster level corrected for multiple comparisons at p<0.05. Maps represent mean activation after air and CO inhalation, and contrasts between protocols. Air>CO (blue-lightblue).

*Visual task (Figure 4).* The flashing checkerboard generated activation in the visual cortex (mean activation) for all scans. Group comparisons showed lower activation in the CO protocol compared to air in the visual cortex, right thalamus and left middle temporal gyrus.

**Figure 4.**
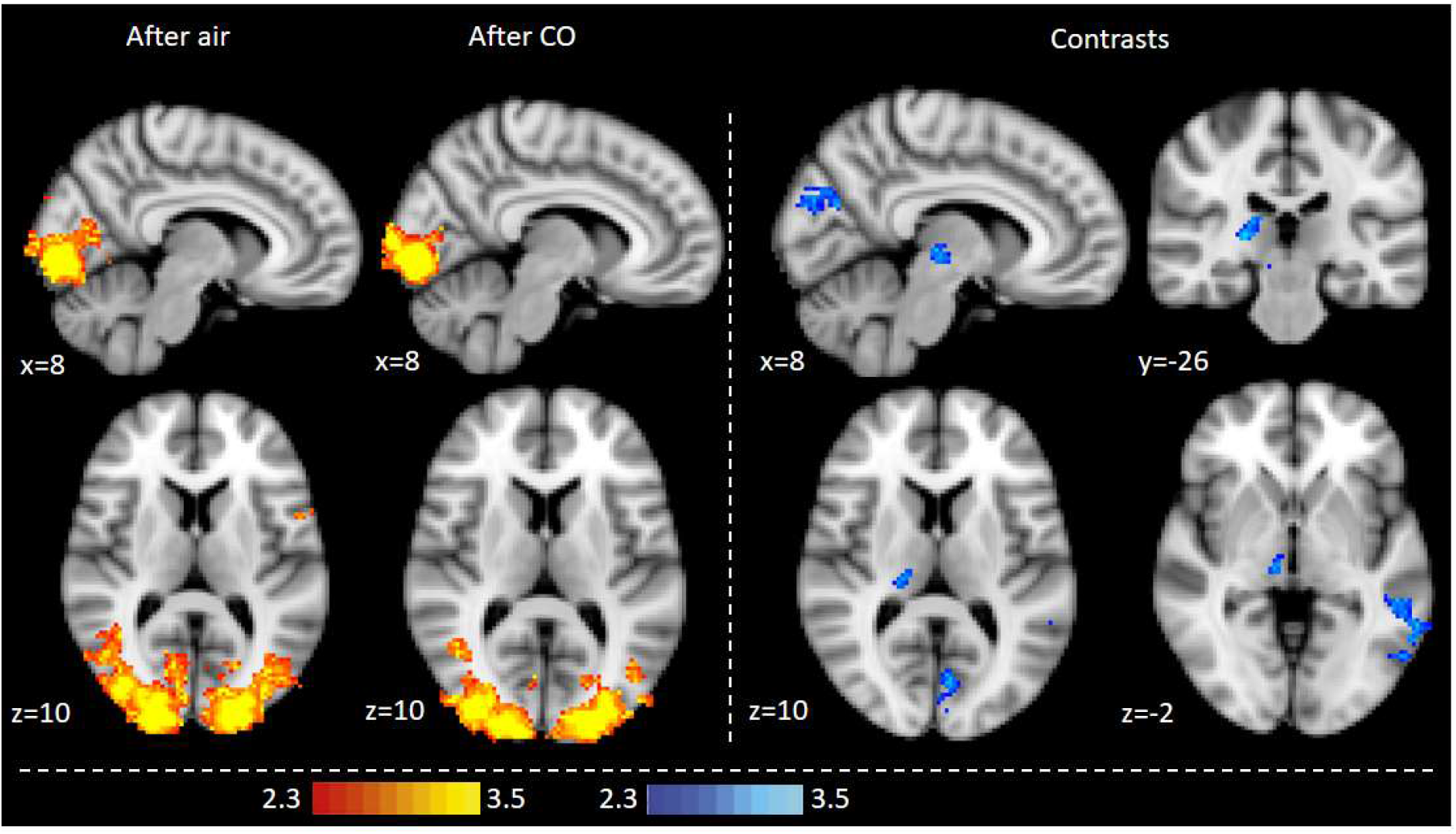
BOLD FMRI activation during visual stimulus. Whole-brain analysis, cluster level corrected for multiple comparisons at p<0.05. Maps represent mean activation after air and CO inhalation, and contrasts between protocols. Air>CO (blue-lightblue).

*Motor task (Figure 5).* The finger-tapping task generated activation in the left primary and secondary somatosensory cortices, the left premotor and primary motor cortices, the left thalamus and the visual cortex for all scans. Group comparisons showed reduced activation in the CO protocol compared to air in the visual cortex and increased activation in the CO protocol compared to air in the premotor cortex.

**Figure 5.**
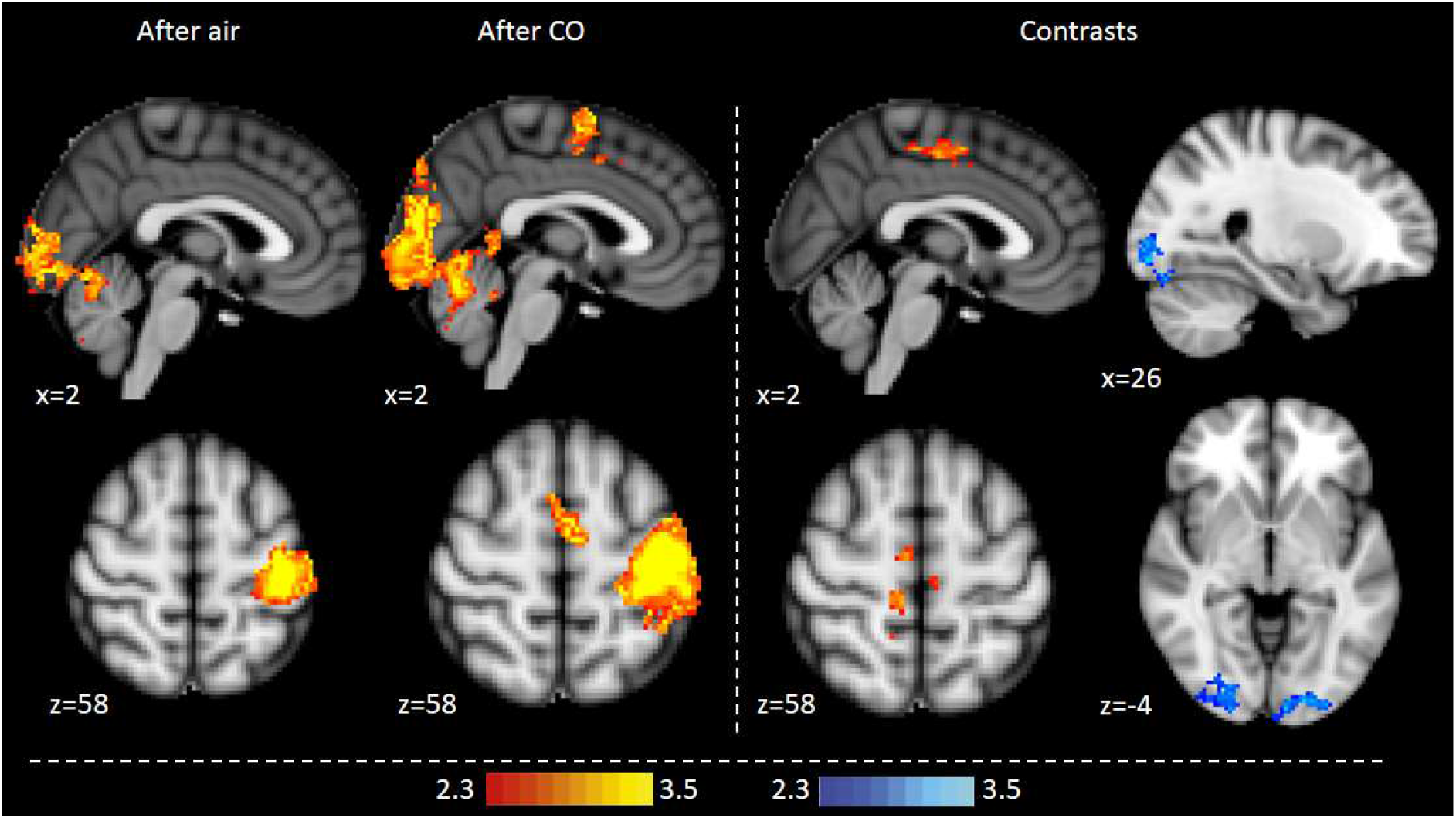
BOLD FMRI activation during motor task. Whole-brain analysis, cluster level corrected for multiple comparisons at p<0.05. Maps represent mean activation after air and CO inhalation, and contrasts between protocols. Air>CO (blue-lightblue); CO>Air (red-yellow).

## DISCUSSION

### Key findings

In this study we show that a small amount of inhaled CO, raising expired levels from 3ppm to 6ppm, significantly alters BOLD signal in never-smokers. This suggests that CO is a non-negligible confound in BOLD FMRI, in particular for studies involving populations regularly exposed to CO (e.g. smokers), even in low doses. Systematic differences in COHb between a patient group consisting of a greater proportion of smokers and a control group of predominantly non-smokers could generate group differences that are CO-related rather than associated with the specific research outcome. This could significantly affect the results of clinical trials and patient-oriented neuroscience research.

### Discussion of findings

Studies have shown that global baseline increases in CBF can reduce task-related BOLD signal as the capacity for further local vasodilation is reduced. Cohen et al. (20) used experimentally induced hypercapnia to reduce visual activation, and Brown et al. (21) showed that the cerebral vasodilator acetazolamide can dampen motor activation. Yet this effect has not yet been linked to CO exposure and smoking. Smoking is associated with a range of diseases, including cardiorespiratory diseases, cancers, dementia and cognitive decline (22) and several mental disorders (23), as well as demographic factors such as socioeconomic status, education and income level (24). CO exposure through cigarette smoking could therefore constitute a significant confound in neuroimaging research.

To probe the vulnerability of the BOLD signal to COHb elevation, we employed a low-level increase in inhaled CO, raising exhaled levels to the lowest associated with tobacco smoking. Using this minimal level, we observed significant effects on BOLD signal during not only a hypercapnic challenge but also commonly used visual and motor tasks. The large impact of low-level CO exposure on common FMRI paradigms such as a simple flashing checkerboard and finger-tapping tasks highlights the relevance of the present findings. Furthermore, the present study consisted of a comparatively small sample of healthy volunteers. Any effects of CO in larger experimental samples are likely to be more robust, further underlining the need to employ care when selecting study populations and recruitment strategies for research involving potentially unbalanced groups of smokers versus non-smokers.

The effect of CO on BOLD signal was not uniform. BOLD signal was reduced after CO exposure in regions associated with breathing and breathlessness during hypercapnia (12, 25, 26), and in the right thalamus, left middle temporal gyrus and visual cortex during the flashing checkerboard (27). However, during the finger-tapping task, CO reduced visual cortex signal (possibly associated with the visual instructions on screen) but unexpectedly also increased motor cortex signal compared to air. This BOLD signal increase appears to contradict the hypothesis that increases in CBF with CO act simply to dampen signal. It has been suggested that hypercapnia may affect BOLD signal differently depending on the type of task and activated brain regions (28). For example, Kastrup et al. (29) reported that BOLD signal changes with hypercapnia were greater in the visual cortex than in the sensorimotor cortex, possibly due to the location of large veins and/or neural activity associated with respiratory stimuli (28). Taken together, this suggests that CO does not have a simple, uniform effect on BOLD signal, but, like hypercapnia, may depend on task and/or region.

### Implications for neuroimaging and clinical trials

In this study, we show that even low-level CO exposure may significantly alter BOLD signal. Due to its affinity for haemoglobin, CO is not readily removed and therefore its effects on signal could persist for some time following inhalation. Here, CO assessments made following the scan (approximately 20 and 30 min after the intervention) show steady, elevated levels of exhaled CO (Figure 2). This level of CO exhalation is at the lower end of that associated with smokers, with mean exhaled values being more than 20ppm in outpatient groups (9). It remains unknown if higher levels of CO exposure will have a greater effect (i.e. a dose-dependent effect similar to that observed in rat aortas (30)). Furthermore, the findings observed in this paper suggest that the effect may be region-or task-dependent, which could complicate any potential adjustments for COHb during analysis. This indicates that CO is an important confound in BOLD FMRI, even in low doses, and its effect sufficiently robust to induce statistically significant changes in even small study populations.

Smoking habit is associated with several diseases, from cardiorespiratory disease and cancer to schizophrenia. Clinical trials in which FMRI is used as an outcome measure may therefore be susceptible to differences in CO exposure. Differences in COHb may occur both longitudinally (if the participants are encouraged to stop smoking as part of their treatment) and whenever patients are compared with controls that are not precisely matched for smoking behaviour. Furthermore, the possibility for dose-dependent effects means that it may not be sufficient to match simply for ‘smoker’ and ‘non-smoker’, but rather the amount of COHb present in the blood stream. Given that only a small increase in COHb may affect BOLD signal, this confound should be monitored carefully, particularly in clinical trials.

### Potential mechanisms

As absolute CBF was not measured in the present study, we cannot be certain that CBF was the cause of the observed dampening of the BOLD response. Indeed, as suggested previously, the mechanism underlying CO mediation of BOLD signal may be more complex. Other mechanisms may contribute to the observed BOLD signal change. For example, the formation of COHb at the expense of oxyhaemoglobin may cause increased CBF through the development of hypoxia (31). While we observed reduced P_ET_O_2_ during the second scan on each experimental day, this was similar for both protocols, and may thus rather be due to altered breathing patterns during the experimental protocol despite pre-scan acclimatization to the breathing system. Furthermore, P_ET_O_2_ remained within normal range throughout the experiment. It is therefore unlikely that hypoxia is the cause of the observed group differences. CO may also slightly inhibit cell respiration even under normoxic conditions (32) and it remains unknown whether the observed effect on BOLD signal is linked in full or part to metabolic modulation. Similarly, we cannot rule out the possibility that CO altered BOLD signal through its role as an endogenous neurotransmitter (33). Participants showed no change in reaction times with CO compared to air, no difference in anxiety scores, and were not able to tell which protocol they were undertaking when prompted. It is thus unlikely that the effect on BOLD signal observed in our study is driven by behavioural factors. While further work is required to elucidate the precise mechanism underlying our findings, it is clear that CO alters BOLD signal and should be considered a non-negligible neuroimaging confound.

### Conclusions

We conclude that even small amounts of inhaled CO can significantly alter BOLD signal during simple tasks such as breath hold, visual and finger-tapping tasks. Further research is required to assess the precise underlying mechanism of this effect. Care should be taken to control for CO as a confound in neuroimaging research when appropriate, for example in studies on clinical populations with greater/lower prevalence of smokers.

## ACKNOWLEDGEMENTS

We would like to thank Steve Knight for his generous assistance with data collection, and Dr Olivia Faull, Dr Anja Hayen and Dr Kyle Pattinson for their invaluable feedback on the analysis and manuscript. This study was funded by the Oxford Brookes University Central Research Fund.

